# Ring shape Golden Ratio multicellular structures are algebraically afforded by asymmetric mitosis and one to one cell adhesion

**DOI:** 10.1101/450528

**Authors:** William E. Butler, T. Bernard Kinane

## Abstract

Golden Ratio proportions are found throughout the world of multicellular organisms but the underlying mechanisms behind their appearance and their adaptive value if any remain unknown. The Golden Ratio is a real-valued number but cell population counts are whole numbered. Binet's formula connects the Golden Ratio to the whole numbered Fibonacci sequence (*f*_*n*+1_ = *f*_*n*_ + *f*_*n*–1_ where *f*_1_ = 1 and *f*_2_ = 2), so we seek a cellular mechanism that yields Fibonacci cell kinetics. Drawing on Fibonacci’s description of growth patterns in rabbits, we develop a matrix model of Fibonacci cell kinetics based on an asymmetric pause between mitoses by daughter cells. We list candidate molecular mechanisms for asymmetric mitosis such as epigenetically asymmetric chromosomal sorting at anaphase due to cytosine-DNA methylation. A collection of Fibonacci-sized cell groups produced each by mitosis needs to assemble into a larger multicellular structure. We find that the mathematics for this assembly are afforded by a simple molecular cell surface configuration where each cell in each group has four cell to cell adhesion slots. Two slots internally cohere a cell group and two adhere to cells in other cell groups. We provide a notation for expressing each cell’s participation in dual Fibonacci recurrence relations. We find that single class of cell to cell adhesion molecules suffices to hold together a large assembly of chained Fibonacci groups having Golden Ratio patterns. Specialized bindings between components of various sizes are not required. Furthermore, the notation describes circumstances where chained Fibonacci-sized cell groups may leave adhesion slots unoccupied unless the chained groups anneal into a ring. This unexpected result suggests a role for Fibonacci cell kinetics in the formation of multicellular ring forms such as hollow and tubular structures. In this analysis, a complex molecular pattern behind asymmetric mitosis coordinates with a simple molecular cell adhesion pattern to generate useful multicellular assemblies. Furthermore, this reductively unifies two of the hypothesized evolutionary steps: multicellularity and cellular eusociality.

## Introduction

The Golden Ratio 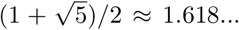 was recognized at least as early as 500 BCE by Phidias, after whom its symbol Φ remains named. Its presence in plants, mollusks, and vertebrates has been commented by naturalists over the centuries, has been depicted in the arts, and has been the subject of teleological conjecture [1, 2]. While it appears in cellular automata[3, 4], the molecular or cellular mechanisms for its presence in multicellular organisms remain unknown. Moreover, it remains unknown how it confers adaptive biological benefit, or if it does at all. We inspect the mathematics of the Golden Ratio for hints as to how and why it appears in multicellular organisms.

The Golden Ratio is a real number but cell population counts are measured by the whole numbers (0, 1, 2, 3, …), so we focus our mathematical search on them. The Fibonacci numbers are the particular whole numbers that obey the recurrence relation *f*_*n*+1_ = *f*_*n*_ + *f*_*n*–1_, where *f*_1_ = 1 and *f*_2_ = 2, giving the Fibonacci sequence (1, 2, 3, 5, 8, …). We employ in this paper the convention common in the combinatorial literature of *f*_1_ = 1 and *f*_2_ = 2 (other bodies of work employ *F*_0_ = 0 and *F*_1_ = 1). Binet’s formula connects the Golden Ratio to the Fibonacci numbers,

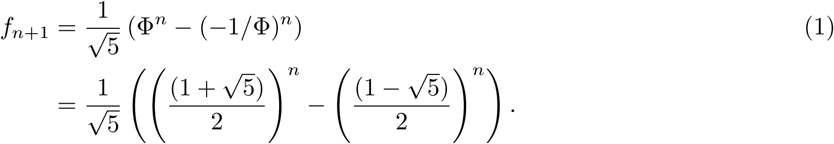

We propose that Golden Ratio patterns in multicellular organisms could be grounded in Fibonacci population kinetics. We propose that the mathematics of Fibonacci cell kinetics are afforded by an asymmetric pause between progeny cells before they undergo mitosis. We propose that candidate molecular mechanisms for kinetically asymmetric mitosis might be found among the described systems for asymmetric epigenetics.

## Dual Combinatorial Engagement

Let us assume that there exists a cell kinetic pathway for a suspension of early progenitor cells to mitose into a population of Fibonacci-sized cell groups. We need furthermore to describe how these might assemble into a larger multicellular structure with Golden Ratio patterns. We propose that the underlying mathematics for this assembly are afforded by a simple molecular cell surface configuration where each cell in each group has four cell to cell adhesion slots.

The Fibonacci numbers partake of a rich set of combinatorial identities [5]. Many of these identities describe how groups of smaller Fibonacci numbers may combine to form larger ones, with the recursive relation *f*_*n* +1_ = *f*_*n*_ + *f*_*n*–1_ being but the simplest. We propose that this configuration of cell to cell adhesion slots should allow Fibonacci-sized cell groups to recombine according to those identities, offering pathways for larger Fibonacci-sized multicellular structures to be assembled from smaller ones.

Four adhesive slots are the minimum for each cell to adhere to two other cells within its Fibonacci-sized cell (to hold the group together) and to adhere to two other cells in two other Fibonacci-sized cell groups. This permits chaining between them. We separate cell to cell adhesion by function into intra and inter Fibonacci-sized group adhesion but the same molecular adhesion machinery may serve both functions since in each case adhesion is between one individual cell and another.

We provide an over and under arrow notation to track the engagement of each cell in two intergroup Fibonacci combinatorial identities. To match the notation by function we classify the vertical slots as intergroup. We classify the horizontal slots primarily as intragroup except at boundaries between groups where they bound two cells from two groups across a gap. We treat the horizontal slots as implied and focus the notation on the vertical ones since they express the binding between Fibonacci-sized groups. The equals sign of an identity is replaced in this notation by an arrowhead and the plus sign by the intersection of arrow lines. The operands to the plus operator are the numbers of cells indicated by the arrow lines’ originating groups. In this simplified treatment we assume there may be multiple operands on the left hand side of an equal side and a single result on the right hand side. We do not allow a cell or cell group to bind to itself.

In this notation, for example, we might represent *f*_1_ + *f*_2_ = *f*_3_ in the top vertical slots as

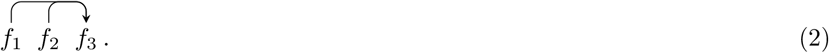

This may represent the biological process whereby the top intergroup cell adhesion slots of cell groups of sizes *f*_1_ and *f*_2_ adhere to and occupy the top adhesion slots of a cell group of size *f*_3_. The vertical slots have size given by the Fibonacci number. This notation is specific to this application and is unrelated to the Fibonacci number of a graph[6]. A triplet as this is the simplest Fibonacci adhesion event involving cell groups of different sizes.

We explore how mitosis might yield Fibonacci-sized groups then we employ this notation to analyze how a nascent population of Fibonacci-sized groups might recombine.

## Asymmetric Mitosis

Binet’s formula implies that Golden Ratio multicellular biofractals might appear if cell population sizes were to grow not as powers of two as governed by classical mitosis (1 → 2 → 4 → 8 → …), but according to the Fibonacci numbers,

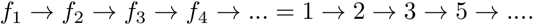

Fibonacci described a rabbit reproduction model based on asymmetric pause between reproduction (Fig 1) [7]. We adapt this model to cell kinetics, substituting the replication unit of a rabbit pair with a cell in a multicellular organism. We assume that cells in a group share an equivalent interval between mitotic cycles. There is an evolving description of the biomolecular infrastructure that might coordinate the timing of mitosis in groups of cells[8, 9, 10]. In the models of asymmetric mitosis below we name cells within a kinetic group with lower case characters.

**Figure 1.**
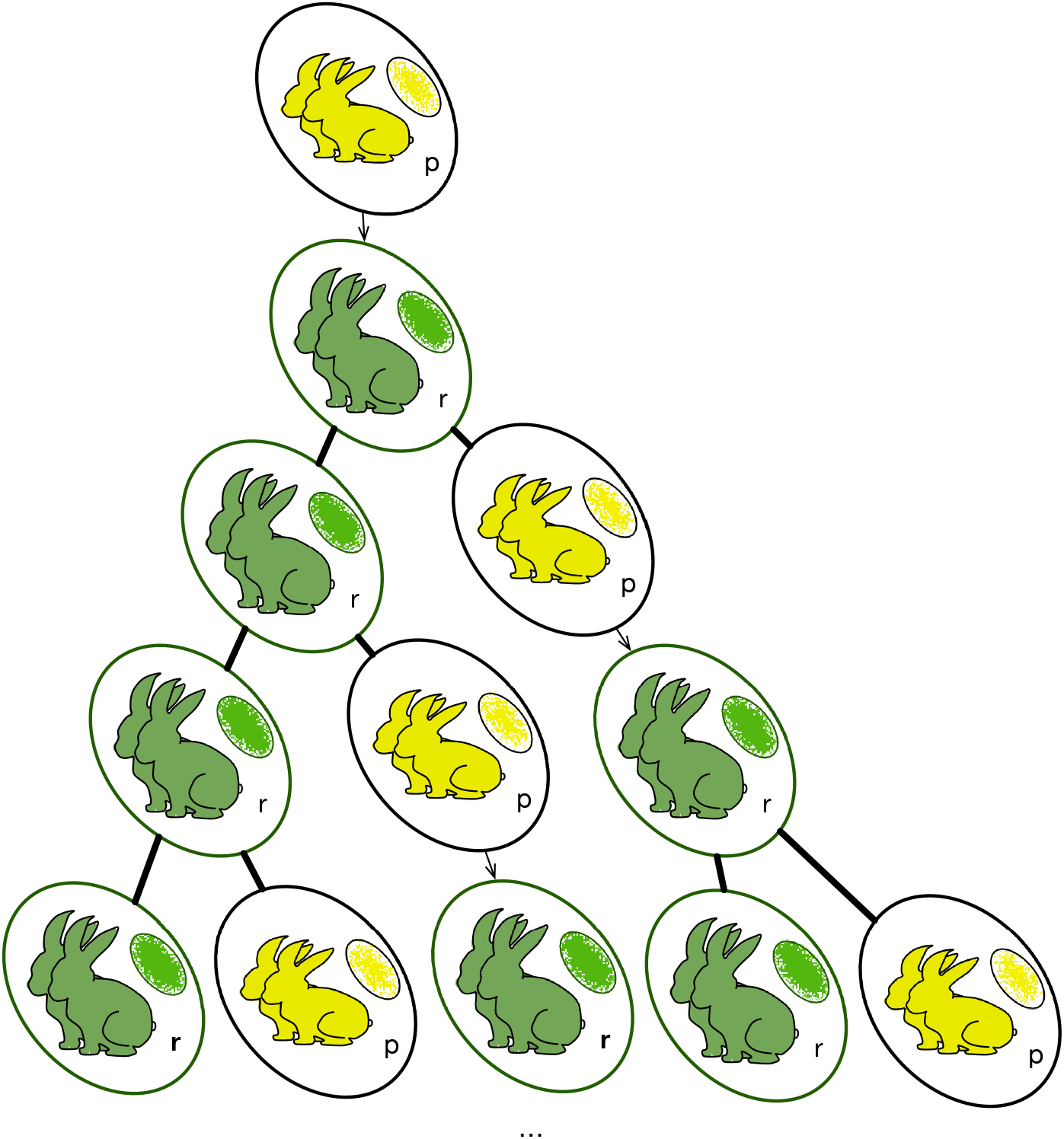
Fibonacci population growth in rabbits and cells with asymmetric pause after each reproduction. Each row is termed a group. The thin horizontal lines represent intragroup adhesions. Similarly, each cell also has unoccupied intergroup cell adhesion slots (not shown).

We consider two cell classes, *r* (replicate) and *p* (pause), of different kinetic properties but otherwise of the same phenotype. In each generation interval an *r* cell replicates to another *r* cell plus a *p* cell, and a *p* does not divide but matures into an *r* cell that divides in the following generation,

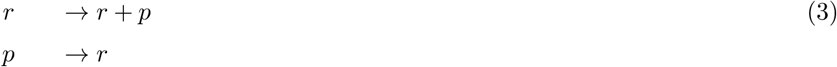

Under Lindenmayer, this corresponds to a propagating, deterministic 0L model[11, 12, 13]. Using subscripting to indicate the number of *r* and *p* cells in a bud at generation *n*, we then have for a generation

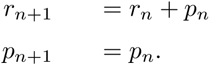

In matrix notation this is

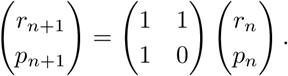

By recursion, the formula for the population at the *n* + 1*^th^* generation is then

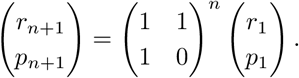

The matrix

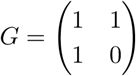

formally connects this model of asymmetric mitosis to the Fibonacci sequence [14, 15].

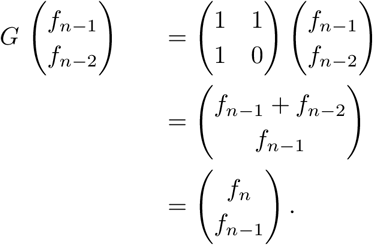

The matrix *G* on its own can produce any three sequential Fibonacci numbers *f*_*n*–2_, *f*_*n*–1_, *f_n_* as[15, 16]

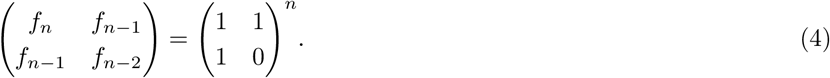

The eigen decomposition of the matrix *G* connects the Fibonacci sequence to the Golden Ratio, Φ, because *G* has eigenvalues λ_1_ = Φ and 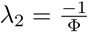 and eigenvectors

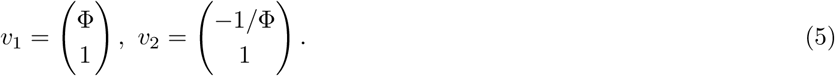

By the definitions of eigenvector and eigenvalue, the structure of *v*_1_ and the value of λ_1_ indicate that if *r_i_* and *p_i_* are in a Golden Ratio, they will both increase by a Golden Ratio in the next generation. This result is consistent with Binet’s formula for calculating an arbitrary Fibonacci number from the Golden Ratio.

## Candidate Mechanisms of Asymmetric Mitosis

Perhaps the molecular machinery controlling the cell cycle is unevenly mixed by chance following telophase often enough so as generally to favor asymmetric mitosis. We reject this as unlikely to be robust.

In the standard paradigm of molecular biology, the mitotic cell cycle includes the dissociation of a double strand of DNA into two single strands at anaphase, each of which forms a template for the synthesis of a complementary strand. Each daughter cell receives one of the two identical copies of double stranded chromosomal DNA, less any DNA replication errors. Meiosis, by contrast, necessarily has genetically asymmetric progeny so its mechanisms of asymmetry are distinct from any contemplated for mitosis.

Perhaps the molecular machinery controlling the cell cycle is unevenly mixed by chance at telophase often enough so as generally to favor asymmetric mitosis. In this model, progeny cell asymmetry is by the actions of random chance on classical mitosis. We reject this as unlikely to be robust.

DNA methylation appears to be an ancient property of eukaryotic genomes that play a role in development in both plants and animals[17, 18, 19, 20] The 5-methylation of cytosine in DNA is a candidate mechanism because at mitotic chromosomal sorting during anaphase only one of the two double stranded DNA daughter strands will be methylated at that position because at replication the complementary guanine will be matched by an unmethylated cytosine (Fig 2) [21]. Cytosine is methylated at the same 5 position that uracil is methylated to give thymine. The steric specificity for methylation at the 5 position of a pyrimidine chemically associates cytosine 5-methylation with the origin of DNA from RNA, but the paleobiochemistry of DNA methylation is an active target of investigation[22, 23].

**Figure 2.**
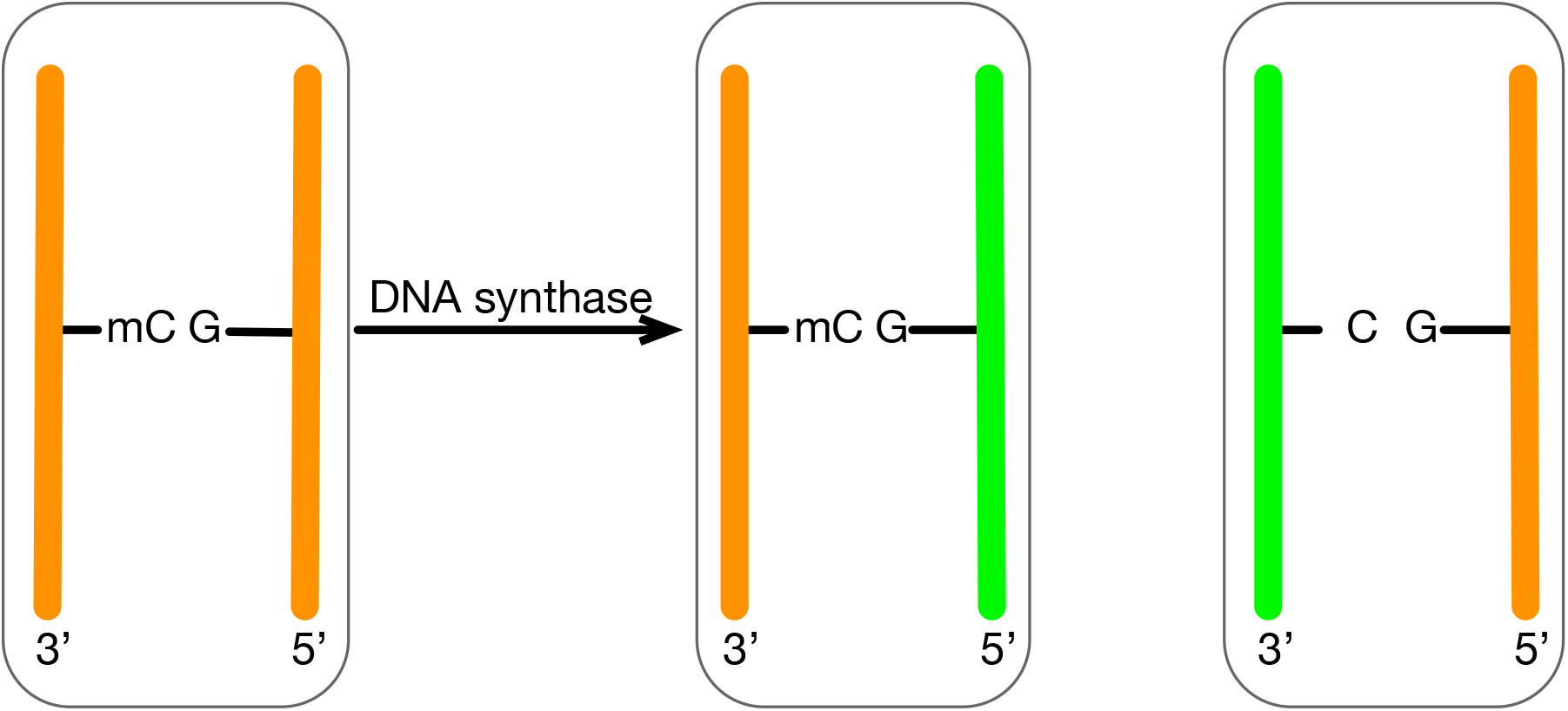
Cytosine methylation creates epigenetic asymmetry of chromosomes in the cell cycle before mitosis.

The histone proteins represent another candidate mechanism for asymmetric epigenetic transmission. They help compact DNA into the confines of a cell, appear subject to regulation including by covalent modification, such as by acetylation and could alter cell cycle regulation [24, 25, 18, 21]. At chromosomal assortment in mitosis, modified histone proteins may assort asymmetrically, and hence be transmitted asymmetrically to the mitotic progeny [24, 26].

The model of Fig 2 depicts one chromosome but an organism may have multiple chromosomes. Regardless of ploidy, candidate mechanisms based on epigenetic asymmetry call for such asymmetry among the chromosomes at anaphase so as to produce Fibonacci growth patterns.

## Recombinatorial Growth

We exclude from consideration reflexive arrangements such as

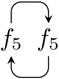

and

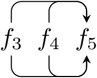

that interrupt the chaining.

With this restriction, the presence of two vertical slots implies a potential for infinite chaining. For example, a set of Fibonacci-sized cell groups *f*_1_, *f*_2_, *f*_3_, *f*_4_, *f*_5_, *f*_6_, … produced by asymmetric mitosis might assemble by overlapping triplet adhesion into a scaffold as

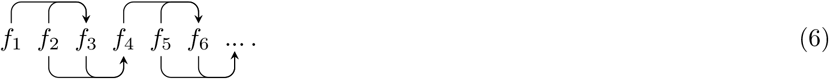

A biological structure of course will not reach infinite size. Yet the capacity for dual Fibonacci engagement implies a path for a multicellular organism to display a self-similar scaffold with Golden Ratio biofractal patterns across a broad range of spatial size. A bio-fractal form may offer the biological efficiency of reuse of the same cell to cell adhesion molecules to maintain structural integrity across spatial scales. Specialized adhesion mechanisms are not required to bind Fibonacci-sized groups of various sizes.

There are Fibonacci identities that produce a larger Fibonacci number from a collection of smaller ones. An example is

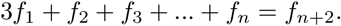

If this represented a collection of Fibonacci-sized groups that adhered as

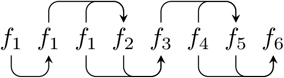

then as per the identity the assembly would have *f*_8_ cells. Suppose a Fibonacci-sized group of cells were to have a way internally to coordinate kinetically asymmetric mitosis mitosis regardless of how it came to be[8, 9, 10]. Then the *f*_8_-sized group might alternately mitose into a *f*_9_-sized group or assemble with other Fibonacci-sized cell groups. An equivalence of mitosis and assembly implies recursive growth.

### Multicellular Rings

Let us assume that a scaffold made of dual Fibonacci combinatorial elements tends to seek a combinatorial arrangement that is fully engaged, meaning where all vertical positions of all symbols participate uniquely in one equation whether on the left hand side as an operand or the right hand side as a result. This corresponds to a biological interpretation where where all cell to cell adhesion slots are occupied. We observe that an open scaffold such in Equation 6 based on concatenated triplets cannot have full engagement at minimum because each end has one vertical slot that is not engaged.

However, if the two ends have an unengaged vertical slot of equal size, then the structure can fold and bind into full engagement. An example is the palindrome

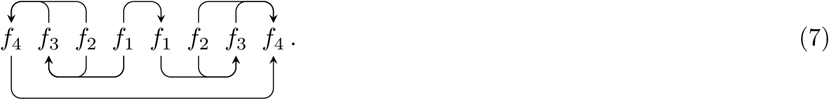

A palindrome is not the only closed form with full engagement. In S1 File we prove a set of conditions where a fully engaged ring can be made from random triplets of Fibonacci groups. With that proof in hand, we generate random numbers that conform to the terms of the proof to guarantee that the result will be a fully engaged scaffold ring (for example see Fig 3).

**Figure 3.**
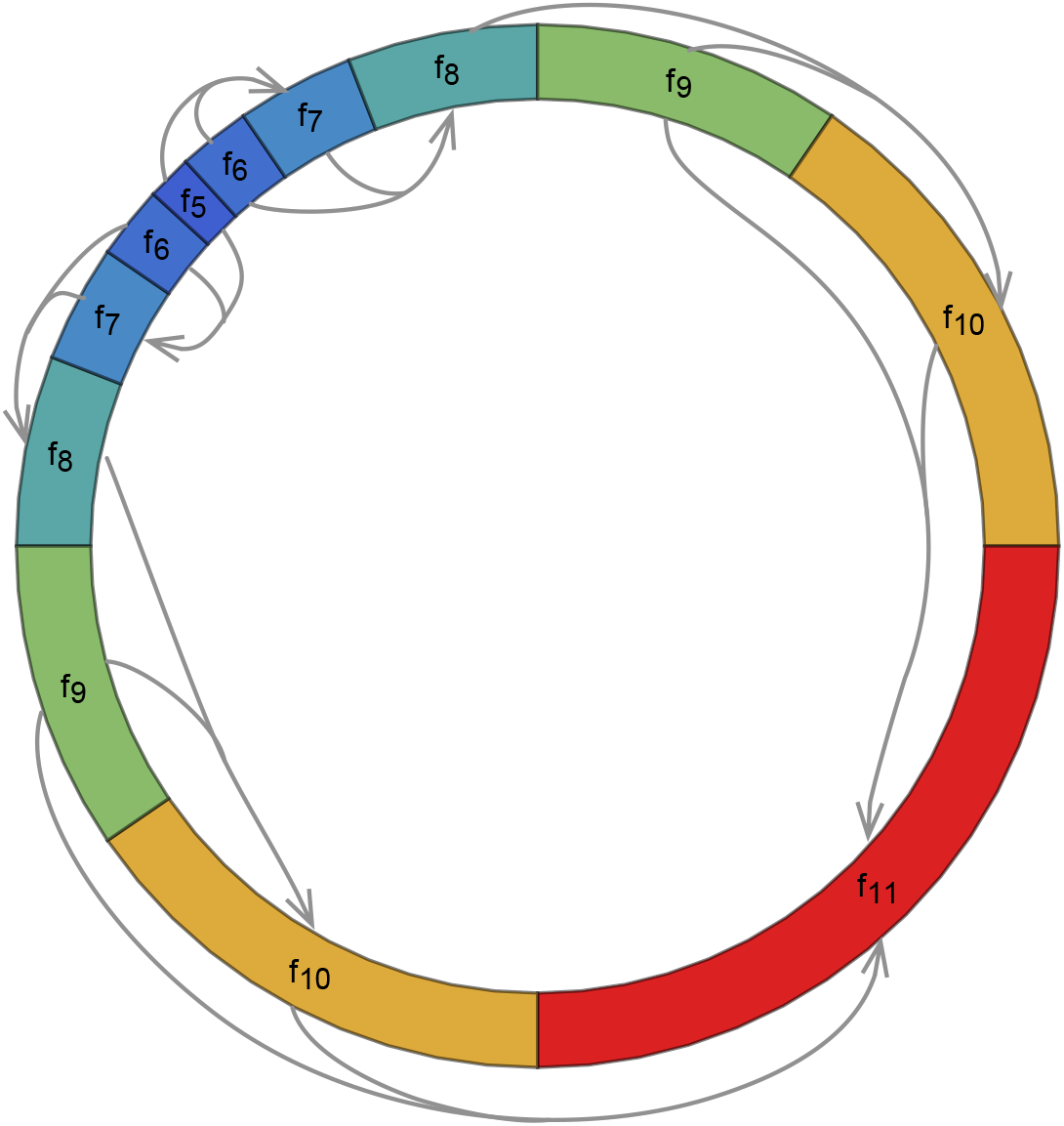
A Scaffold Ring. All inter and intragroup adhesion slots are occupied. The arrows depict the intergroup adhesion slot bindings.

If a number of ring shapes of the same diameter were to stack up above and below the plane of the page and each cell were to have a further pair yet of unoccupied intergroup adhesion slots oriented in the third dimension perpendicular to the plane of the page, then the rings could anneal into a tubular structure. We use here the Fibonacci numbers to count the number of cells in a cell group, but the Fibonacci numbers offer a different combinatorial interpretation. For an open form of length *n* cell diameters, the Fibonacci number *f*_*n*_ carries the combinatorial interpretation of counting the number of ways to fill those *n* cell diameters with single cells and cell pairs. For a ring with circumference *n* cell diameters, that count is given by the Lucas number *L_n_*[5, 16, p 17]. For a given number of cell diameters, the Lucas numbers are larger than the Fibonacci numbers, as per the identity

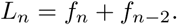

Since for the same number of cells the Lucas number is larger than the Fibonacci number, a ring shape as a template offers a larger number of ways for single cells and pair cells to anneal to it than does a non-branching open shape. As with the Fibonacci numbers, the ratio between adjacent Lucas numbers tends toward the Golden Ratio.

## Individual Replicative Restraint Algebraically Promotes Fractal Collaboration

In classical mitosis, cell populations grow as powers of two (1, 2, 4, 8, …) but in kinetically asymmetric mitosis as modeled here, cell populations grow as the Fibonacci numbers (1, 2, 3, 5, …). All else being equal, classical mitosis enjoys a reproductive advantage since 2*^n^* grows faster than *f*_*n*_ for increasing *n*. Kinetically asymmetric mitosis requires the further expenditure of a relatively complex asymmetric molecular subsystem such as asymmetric epigenetics.

However, equipped with a molecularly simple cell to cell adhesion tendency, individuals in these cell groups may engage in dual Fibonacci combinatorial identities with individuals in other cell groups to assemble efficient multicellular biofractals inclusive of rings. It remains unknown how multicellular organisms develop luminal specializations, but they might represent an extension into three dimensions of the process described here. Insofar as kinetically asymmetric mitosis produces one reproductive and one transiently non-reproductive “caste”, this analysis algebraically unifies two hypothesized major evolutionary transitions: the formation of multicellular forms and of eusocial organizations[27, 28, 29]. The Golden Ratio arises less as a universal constant but more as an spatial side effect of coordinated mitosis and assembly[30].

This notation expresses the formation of multicellular rings as an algebraic possibility but no argument is made as to thermodynamic probability. Further research is required to characterize the potential impacts of statistical errors in the processes and of deviations in the molecular mechanisms. We emphasize the analysis of triplets, but many other forms may be considered. This analysis could be furthered by the incorporation of work on cellular automata[12, 31, 32, 33]. Further research is required to analyze three-dimensional configurations of Fibonacci-sized cell groups, their combinatorial assembly properties, and their thermodynamic constraints.

## Supporting information

## Author’s contributions

Both authors contributed equally.

## Acknowledgements

Elizabeth G. Shannon and Ziv Williams commented on versions of the manuscript.

## Additional Files

Additional file 1 — Sample additional file title

Random combinatorial scaffold rings.

