## Supplementary material for "Ring shape Golden Ratio multicellular structures are algebraically afforded by asymmetric mitosis and one to one cell adhesion"

### 1 Random Ring Patterns

To nourish our intuition we wish to generate a family of random scaffold rings having full adhesion engagement. To start, we offer some definitions and provide a theorem showing that as scaffold ring as defined here has full adhesion engagement.

**Definition 1** *A Fibonacci triplet is a set of three groups of sequential Fibonacci size that have intergroup receptor bindings to each other. A relative up triplet has increasing Fibonacci numbers such that a relative up triplet to* $f_{k-1}$ *is*

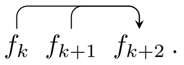


*and a relative down triplet is*

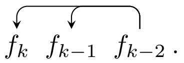


It does not matter in this definition whether the intergroup receptors are top or bottom.

**Definition 2** *A triplet scaffold is a structure that is composed of concatenated Fibonacci triplets where all slots are engaged and the sumands and sums are contiguous.*

We observe that in such a structure, a given triplet has six vertical adhesion slots. Two serve as sumands to bind the concatenated triplet to the right. One serves as a sum to bind to two sumands from the triple to the left. These overlapping Fibonacci relations bind the triplets one to the next in the scaffold. Three slots are internal to the triplet. Two of those are summands for a sum in a third slot.

**Definition 3** *A scaffold ring is a triplet scaffold where the ends have folded to meet and adhere. An example is*

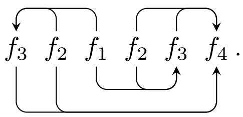

Start with an arbitrary Fibonacci number, $f_k$, append to it an equal number of relative up and down triplets in arbitrary order, and adhere the slots.

**Theorem 1** *Such a ring has full adhesion engagement.*

*Proof* Start with an arbitrary initial Fibonacci number $f_k$. Append a relative up triplet. The rightmost group is of size $f_{k+3}$ because appending a triplet adds 3 to the rightmost Fibonacci index.

$$f_k \quad f_{k+1} \quad f_{k+2} \quad f_{k+3} \;.$$

Append a relative down triplet. This subtracts 3 from the rightmost index.

$$f_k \quad f_{k+1} \quad f_{k+2} \quad f_{k+3} \quad f_{k+2} \quad f_{k+1} \quad f_k \;. \tag{1}$$

There are 5 internal Fibonacci-sized groups. These have 10 adhesion slots available. There are 6 sumand and 4 sum slots occupied, so the internal groups have full engagement. The groups on the left and right ends each have $f_k$ occupied and $f_k$ unoccupied slots. If the structure folds, the $f_k$-sized slots on each end will adhere because they are o the same size. Between the two triplets we can insert an equal number of relative up and down triplets in random order and there will still be adhesion closure because each end will have $f_k$ unoccupied vertical adhesion slots.

In the paragraph above, we start with a relative up triplet and end with a relative down triplet. We find however that the same argument holds whether we start or end with an up or down triplet so long as the number of them is equal. We also find that the order of the relative up and down triplets does not matter.

□

**Lemma 1** *In a fully engaged ring, duplicating a Fibonacci-sized group number or replacing a duplicate with a single produces a longer or shorter ring that still has full engagement.*

This is because inserting a duplicate number only flips the vertical polarity of the ensuing receptor orientation. This lemma is nonsensical in the case of a palindromic ring where its successive application would cause the removal of all groups. Hybrid forms with an inner loop and open ends are algebraically possible but not treated here.

Removing the duplicate $f_k$ from Equation 1 produces

$$f_{k+1}\ f_{k+2}\ f_{k+3}\ f_{k+2}\ f_{k+1}\ f_k\,.$$

This is a triplet scaffold where the adhesions are fully engaged.

With this proof and lemma we program a computer to generate some random scaffold rings with full engagement. We randomly generate a set up relative up and down triplets, randomly insert and remove duplicates, and graph some results in Fig 1 to portray a range of allowed woven rings.

### Figures

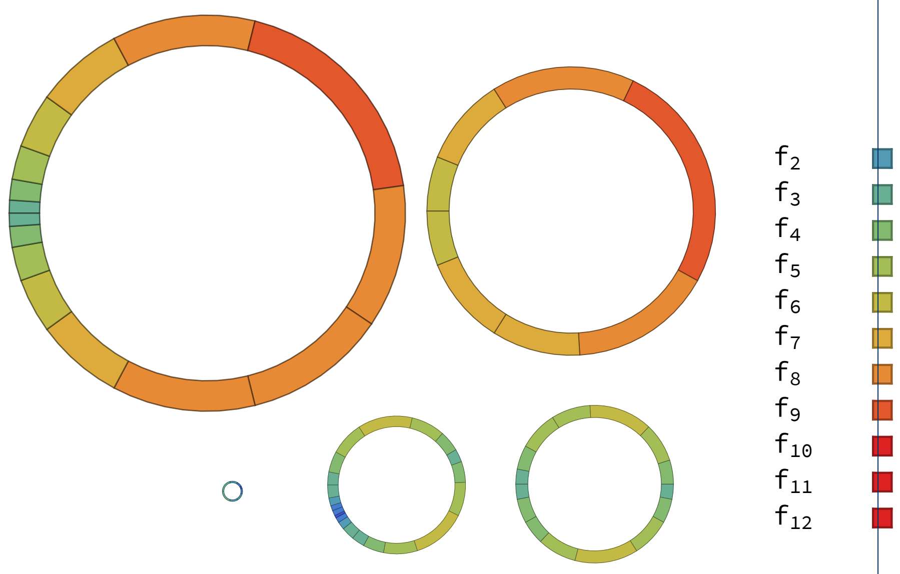


**Figure 1 Some Random Woven Rings.** The legend gives the Fibonacci color code scheme. Receptor relationships are not depicted.
